# H2B.W2, a Spermatocytes-specific Histone Variant, disrupts nucleosome stability and reduces chromatin compaction

**DOI:** 10.1101/2024.12.12.626759

**Authors:** Thi Thuy Nguyen, Dongbo Ding, Xingpeng Bai, Matthew Y.H. Pang, Mingxi Deng, Yue Liu, Tingyu Jin, Zhichun Xu, Yingyi Zhang, Yuanliang Zhai, Yan Yan, Toyotaka Ishibashi

## Abstract

Spermatogenesis is a highly regulated process that requires precise chromatin remodeling, which includes the incorporation of testis-specific histone variants. While several of these variants have been characterized, the role of H2B.W2, a member of the H2BW family, remains largely unclear. Here, we showed that H2B.W2 expression occurs mainly in spermatocytes, slightly later than its paralog H2B.W1. Cryo-electron microscopy (cryo-EM) analysis of H2B.W2-containing nucleosomes reveals a more relaxed conformation compared to canonical nucleosomes caused by weakened interactions between the outer DNA turn and the histone core. We pinpointed the N-terminal tail and α2 helix of H2B.W2, specifically residues D85 and Q101, as critical for nucleosome destabilization. Furthermore, we identify G73 within the L1 loop as a key residue involved in disrupting higher-order chromatin structure. Our findings suggest that H2B.W2-mediated nucleosome and chromatin destabilization may play a role in regulating gene expression during spermatogenesis, with potential implications for sperm development and function.

## Introduction

Spermatogenesis is the process in which mature spermatozoa develop from primordial germ cells in the testis. During this process, chromatin in the germ cells and their progenies undergoes a series of critical changes, including histone acetylation and phosphorylation, the incorporation of testis-specific histone variants, and the transition of histones to protamines as round spermatids develop into elongated spermatozoa(1). The incorporation of testis-specific histone variants plays important roles in spermatogenesis by regulating chromatin structure and gene expression.

Histone proteins are fundamental components of the chromatin structure. In addition to the canonical H2A, H2B, H3 and H4 histones, there is a diverse group of histone variants. Many of these variants, including H2A.B (H2A.Bbd), H2B.C1 (TH2B), H2B.W1 (H2BFWT), H3.4 (H3T), and H3.5, are specifically expressed in the testis. Multiple testis-specific histone variants are known to be key regulators of gene expression and chromatin structure during specific spermatogenic events. For example, H3.4 is essential for entry into meiosis, and knock-out mice lacking H3.4 fail to enter meiosis(2). H2B.C1 is first expressed in spermatocytes and replaces the bulk of the canonical H2B in mature spermatozoa(3). Disruption of both *H2ac1 (TH2A)* and *H2bc1*, causes spermatogenesis defects in mice(3,4).

In addition to H3.4 and H2B.C1, H2B.W1, which is an H2BW family histone variant only found in primates, is only expressed in the mid-to-late spermatogonia during spermatogenesis(5). It has an extended N-terminal tail, which prevents it from participating in mitotic chromosome assembly(6,7). GFP-H2B.W1 was localized at telomeric region in the Chinese hamster lung V79 cell line(6). Single nucleotide polymorphisms of the *H2BW1* at - 9C>T (which causes a frameshift termination) or 368A>G (which alters H100>R) are associated with male infertility in humans(8,9). The H2B.W1 nucleosome is destabilized, and the instability of the nucleosome is further exacerbated by the H100R mutation(5), which is link to male infertility.

In contrast to the primate-specific H2B.W1, H2B.W2 (also known as H2B.W histone 2 and H2BFM) is found in multiple mammalian lineages and exhibits a distinct tissue expression, being present in both testis and ovary(10). Such expression profile implies a unique role for H2B.W2 in germ cell development beyond spermatogenesis. Notably, the *H2BW2* gene, located on chromosome X (NC_000023.11 (104039956…104042454) in the human genome, encodes a protein with low sequence conservation across species. H2B.W2 shares only 35% and 71% amino acid identity with canonical H2B and H2B.W1, respectively. This divergence in primary amino acid sequence further supports the idea that H2B.W2 may have distinct functions compared to its homologs. However, the structural and functional properties of H2B.W2, specifically its role in nucleosome stability and chromatin compaction during spermatogenesis, remain largely unexplored.

Here, we characterized the human H2B.W2 nucleosome structure and examined its effect on chromatin conformation. We showed that the bulk of H2B.W2 protein is expressed from late spermatogonia to leptotene spermatocytes, which is slightly later than its closest homolog H2B.W1. Our structural and biophysical analyses revealed that H2B.W2 destabilizes the nucleosome core particle by reducing the energy barrier for the unwrapping of the outer turn nucleosomal DNA by ∼35%. We identified the extended N-terminal tail, and D85 and Q101 in the α2 helix region, as critical for the H2B.W2-induced nucleosome destabilization. Furthermore, we found that the G73 residue in the L1 loop is crucial for the higher-order chromatin compaction. Given the presence of the series of tightly regulated chromatin remodeling processes in spermatogenesis, our findings provide a foundation for understanding the unique role of H2B.W2 in spermatogenesis and its potential implications for proper sperm development.

## Material and methods

### Human testis immunohistochemistry and immunofluorescence staining

Formalin-fixed paraffin-embedded (FFPE) from the non-tumor region of patients with testicular cancer (Origene) were sectioned to a thickness of 8 µm. The sectioned samples were then deparaffinized, rehydrated, and stained with a rabbit anti-*hs*H2B.W2 antibody (prepared in-house, at 1:200 dilution, based on a peptide epitope, 40-54, CRGSRRRHANRRGDS, Shanghai Youke Biotechnology), a rabbit anti-*hs*H2B.C1 antibody (prepared in-house and used at 1:1000 dilution, 2–17, EVSSKGATISKKGFKK, Shanghai Youke Biotechnology), a rabbit anti-H2B.W1 antibody (Abcam, ab185682, at 1:35 dilution), and a rabbit anti-H4 antibody (Abcam, ab7311, at 1:1000 dilution), respectively. A rabbit specific HRP/DAB (ABC) detection IHC kit (Abcam, ab64261) was used for visualization according to the manufacturer’s protocol. Modified Mayer’s hematoxylin (Abcam, ab245880) was used to counterstain the nuclei and the images of the stained sections were captured using an Axio Scan.Z1 slide scanner (ZEISS). Scan.Z1 slide scanner (ZEISS).

For immunofluorescence staining, the sections were co-staining of γ-H2A.X and H2B.W2. A monoclonal mouse anti-phospho-histone H2A.X (Ser139) antibody, clone JBW301 (Sigma-Aldrich, 05-636, at 1:50 dilution) was added to the anti-H2B.W1 antibody solution. After washing in PBST, the sections were incubated with a mixture of Alexa Fluor™ 647 goat-anti-rabbit antibodies (Invitrogen, A-21245, at 1:250 dilution) and Alexa Fluor™488 donkey-anti-mouse (Invitrogen, A-21202, at 1:250 dilution). The cell nuclei were counterstained with freshly diluted Hoechst 33342. Images of the immunofluorescence-stained cells were captured using an LSM 980 (ZEISS) confocal microscope.

### Nucleosome and tri-nucleosome reconstitution

The histone octamer was prepared by mixing an equal molar amounts of each histone H2A, H2B or H2B.W2 (modified H2B W2s), H3 and H4 in 40 μl of unfolding buffer (6 M guanidinium chloride, 20 mM Tris-HCl pH 7.5, 4 mM dithiothreitol), followed by 4 h dialysis against water and overnight equilibrium dialysis in octamer buffer (2 M NaCl, 50 mM Tris-HCl pH 7.5, 0.5 mM EDTA, 5 mM β-mercaptoethanol) at 4 ℃. H2B, H2B.W2, domain swapping H2B.W2 or point mutated H2B.W2 octamers were mixed with 147 bp (for the cryo-EM structure), 184 bp (for the MNase assay), 258 bp (for the salt stability assay), or 507 bp (for tri-nucleosome FRET assay) DNA containing the 601 Widom sequence at an optimized molar ratio in the 2 M reconstitution buffer (2 M NaCl, 10 mM Tris-HCl pH 7.5, 0.5 mM EDTA, 5 mM β-mercaptoethanol), then were dialyzed overnight by gradually decreasing the salt concentration to 0 M. Nucleosome samples were then verified using a 5% polyacrylamide native 1xTBE PAGE.

### Cryo-EM sample preparation

H2B.W2 nucleosome samples were loaded with a 147 bp 601 DNA template, and then gradually cross-linked in the 0 M reconstitution buffer (10 mM Tris-HCl pH 7.5, 0.5 mM EDTA), 0.15% glutaraldehyde (Sigma) at 4°C for 2.5 h. The nucleosome samples were purified using a Mini Prep Cell system (Biorad), after which they were ready to be prepared for cryo-EM preparation. First, the quality of the sample was checked by negative staining using uranyl acetate in a 3.05 mm diameter carbon film copper specimen grid. Images of the stained particles were captured by scanning the specimen grid through a Talos120 kV TEM microscope (HKUST Biological Cryo-EM Center). Those nucleosomes that passed the quality control were then loaded into a freshly glow discharged holey carbon golden grid (Quantifoil™ R 2/2 on 300 gold mesh) (2.5 µg of nucleosome diluted in 40 mM NaCl reconstitution buffer in 3 μl was loaded in the grid). Micrographs of the nucleosome particles were collected with Krios G3i cryo-TEM microscope (Thermo Fisher Scientific) and a K3 camera (Gatan) in the HKUST Biological Cryo-EM Center with 81 000 times magnification at 300 kV. Electron exposure setting was 50 e^−^/Å^2^ and each pixel is 1.06 Å. Other details of the microscope setting details are listed in Table S1.

### Image processing, model building and refinement

Movies were imported into cryoSPARC (version: V4.1.2). Patch motion correction (multi), followed by patch CTF estimation (multi), was performed with default settings. Bad images were removed with the “manually curate exposure” function. One hundred micrographs were used for blob-picker to create a template for the template-picker. An extraction box size was set at 200 pixels for the extract-from-micrographs function. After 2D selection, ab initio reconstruction was performed for 3D classification. Homogeneous refinement (new!) was used for mapping.

The template model (PDB: 3LZ0) was fitted with the density map using Chimera (Version 1.16) and further rebuilt with WinCoot following the developer’s instructions. The model was further refined and validated with the Phenix (version 1.20-4459) real-space refinement function and comprehensive validation (cryo-EM) function.

### MNase assay

The nucleosomes with the 184-bp Widom 601 sequence DNA (2.0 μg for DNA) were mixed with 0.02 unit/ul MNase (Worthington) in 10 µl reaction solution (50 mM Tris-HCl pH 7.5, 40 mM NaCl, 2.5 mM CaCl_2_, and 1.9 mM DTT) and incubated at 37°C for 0, 3, 9, 15 and 21 min. After the incubation, the reactions were quenched by adding 10 µl proteinase digestion buffer (20 mM EDTA, 0.5 mg/ml Proteinase K (Invitrogen 100005393), 20 mM Tris-HCl pH 7.5, and 0.25% SDS). The resulting products were analyzed by non-denaturing 12% PAGE in 0.5x TBE buffer (45 mM Tris base, 45 mM boric acid and 1 mM EDTA).

### Salt stability assay

Equal amounts of nucleosome samples were mixed with different concentrations of NaCl buffer (10 mM Tris-HCl pH 7.5, and 0.5 mM EDTA containing 0, 0.8, 1.6, or 2.4 M NaCl) at a 1:1 ratio v/v, and then incubated for 5 min at room temperature. The samples were then dialyzed in 0 M buffer (10 mM Tris-HCl pH 7.5, and 0.5 mM EDTA) for 15 min, after which they were loaded into a 6% 1xTBE PAGE.

### Single-molecule optical tweezers assay

Single-molecule optical tweezers measurements were conducted on nucleosomes using methods described previously(11,12). The wrapped rate and unwrapped rate (*k_w_* and *k_u_*) were extracted from the inverse of the lifetimes of the nucleosome unwrapped state *t_u-w_* and wrapped state *t_w-u_*, respectively. The equilibrium force (*F_eq_*) and equilibrium rate (*k_eq_*) were defined from the same length of time between *t_w-u_* and *t_u-w_*. The energy barrier of the outer wrap at 0 pN (*ΔG_0_*) was extracted by methods described previously(13–15).

### Fluorescence resonance energy transfer (FRET) assay

To investigate the impact of H2B.W2 on chromatin compaction, we reconstituted tri-nucleosome arrays using ATTO488 and ATTO594 double-end labeled N3-Widom 601-507bp DNA templates(16). These templates are specifically designed for tri-nucleosome reconstitution and contain three copies of the Widom 601 positioning sequence separated two 23-bp linkers, facilitating uniform nucleosome assembly. Following reconstitution, the arrays were dialyzed against 0 M FRET buffer (10 mM Tris-HCl pH 7.5, 0.1 mM EDTA, 5 mM β-ME) for 4 h at 4°C and then concentrated to 200 ng/μl using Amicon Ultra-0.5 centrifugal filter units (Merck). Nucleosomes are then diluted with ice-cold 0 M FRET buffer to a concentration of 50 nM, and adding varying concentrations of MgCl_2_ (0, 0.2, 0.4, 0.6, 0.8, 1.0, 1.6, 2.0, 3.2 mM). After the incubation on ice for 5 min, 40 μl aliquots of each reaction mixture were transferred to a black, flat-bottom 384-well high-throughput plate (SPL Life Sciences). The plate was incubated at 37°C for 15 min, and fluorescence signals were measured using a FlexStation 3 multi-mode microplate reader (Molecular Devices) (excitation/emission wavelengths: ATTO488, 488/515 nm; ATTO594, 594/610 nm). The FRET proximity ratio (P) was calculated using the following equation from Gansen et al.(17):

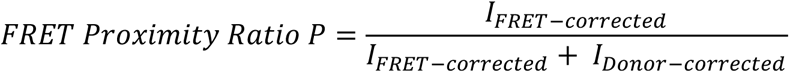

*I_FRET-corrected_* : fluorescence intensity of the sample in the FRET channel after subtraction of fluorescence intensity of the buffer only in the FRET channel

*I_Doner-corrected_* : fluorescence intensity of the sample in the donor channel after subtraction of fluorescence intensity of the buffer only in the donor channel

### Fluorescence recovery after photobleaching (FRAP)

FRAP analysis was conducted on transiently transfected HeLa cell lines expressing histone-E-GFP fusion proteins according to a method from(5). Briefly, HeLa cells were transfected with H2B-GFP or H2B.W2-GFP for 48 h, after which regions of the nucleus were irradiated using a 488 nm laser for bleaching. The fluorescence signal was recorded before bleaching and then at 30-s intervals after bleaching for a total of 40 min.

## Results

### H2B.W2 Expression is Restricted to Early Spermatogenesis

Understanding the temporal and spatial expression patterns of histone variants provides valuable insights into their potential functions during spermatogenesis. Given that the sequence of H2B.W2 is quite different from canonical H2B and another H2BW family histone variant, H2B.W1 (Supplementary Figure 1), we investigated the expression of H2B.W2 in human testis using immunohistochemistry staining with a home-raised anti-H2B.W2 antibody (Figure 1A, and Supplementary Figure 1 and 2). The results showed that H2B.W2 is only expressed in the early stages of spermatogenesis. In contrast, H2B.C1 is deposited on the chromatin of male germ cells during late spermatogenesis (Figure 1A). To determine the H2B.W2 expression pattern, we performed a co-staining of H2B.W2 with phosphorylated H2A.X (γ-H2A.X) (Figure 1B and Supplementary Figure 3). γ-H2A.X patterns can be used to identify spermatogenesis stages with decent accuracy(18,19). Several small DSBs and the γ-H2A.X signal appears in the middle of spermatogonia (ii, iii). The green γ-H2A.X signal is primarily limited in the sex bodies in pachytene spermatocytes (iv), and the signals largely disappeared in a round spermatid (v)(19). Based on the γ-H2A.X signal pattern, we found that H2B.W2 is mainly present in spermatocytes and is markedly decreased in round spermatids. Furthermore, our analysis of a previously published single-cell RNA sequencing dataset further supports these results (Figure 1C). This dataset revealed the distinct expression patterns of several histone variants. H2B.C1 is expressed in the later stages of spermatogenesis, while H2B.W1 is primarily expressed in spermatogonia and leptotene spermatocytes. In contrast, H2B.W2 is expressed slightly later than H2B.W1 in differentiated spermatogonia and spermatocytes(20) (GEO: GSE106487) (Figure 1C and Supplementary Figure 4). Interestingly, H2B.W2 is also expressed in around 40% of H2B.W1 expressing cells, and the expression of H2B.W2 also partially overlaps with that of H2A.X, H3.3, and H3.4 (Figure 1C). Our results indicate that H2B.W2 expression begins in late spermatogonia, then peaked in Leptotene and Zygotene spermatocytes, and decreases in round spermatids (Figure 1D).

**Figure 1.**
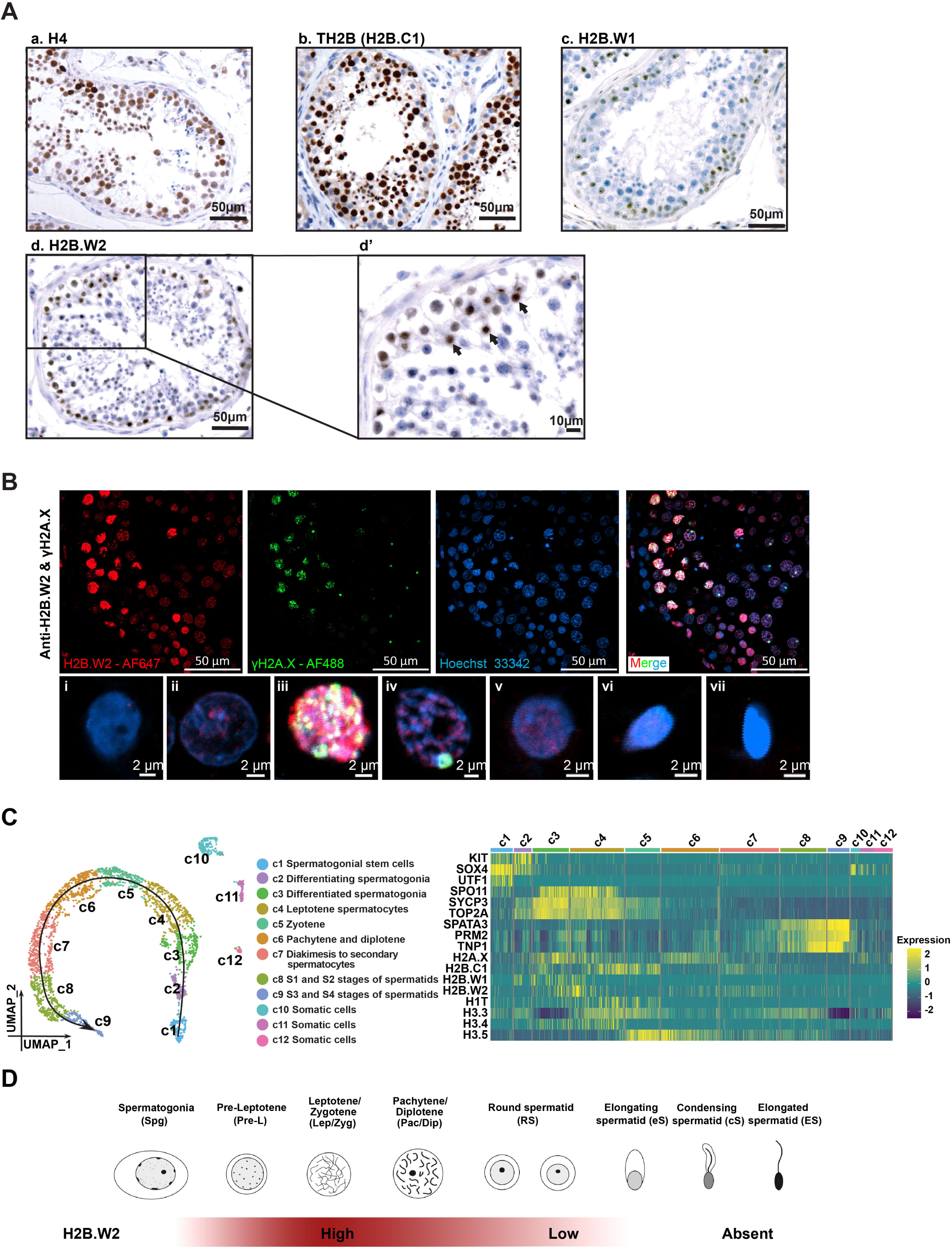
H2B.W2 is mainly expressed in spermatocytes, which is later than another histone variant H2B.W1. (**A**) Sections of human testis were immunostained with an (a) anti-H4, (b) anti-H2B.C1, (c) anti-H2B.W1, or (d) anti-H2B.W2 antibody. The region bounded by the black rectangle in (d) is shown at higher magnification in (d’), where H2B.W2-positive spermatocytes are indicated by black arrows. Scale bars are 50 µm (a-d) and 10 µm (d’). (**B**) (upper panel) Representative immunofluorescence staining examples of human testis sections with anti-H2B.W2 (red) and anti-phospho-histone H2A.X (γ-H2A.X; green) antibodies. The DNA was counter-stained with Hoechst 33342 (blue). Scale bars = 50 μm. (lower panel) Enlarged images of representative cells at different stages of spermatogenesis, which were extracted from the merged image shown in the upper row panel or merged images of samples treated with the same immunofluorescence staining. These images show: an early spermatogonia located at the outermost part of seminiferous tubule without any γ-H2A.X signal in its oval-shaped nucleus (i); a mid-late spermatogonia with very weak γ-H2A.X signal and weak H2B.W2 signal that appear as numerous small foci (ii); an early spermatocyte with strong and globally distributed γ-H2A.X signal and strong H2B.W2 signal (iii); a pachytene spermatocyte with strong γ-H2A.X-staining in the sex bodies and weaker H2B.W2 signal (iv); a round spermatid without γ-H2A.X staining and weak H2B.W2 signal (v); an elongating spermatid with condensed nuclei and almost disappeared H2B.W2 signal (vi); an elongated spermatid with conically-shaped condensed nuclei (vii). H2B.W2 (red), γ-H2A.X (green), and DNA (blue). Scale bars = 2 μm. (**C**) UMAP (Uniform Manifold Approximation and Projection) plots of 2,854 human testicular cells (left panel). This plot is the same as the figure from (Ding et al., 2024) (5). The expression heat map of stage marker genes and histone variants according to different stages (right panel). The single cell RNA sequencing data of human testicular cells were rebuilt from a data set previously published by Wang M. et al., ((20)). (**D**) Schematic view of H2B.W2 expression stages in spermatogenesis.

### H2B.W2 Destabilizes Nucleosome Structure

While this specific expression pattern suggests a potential role for H2B.W2 during meiosis, its precise function at the molecular level remains unclear, especially on the nucleosome level. To gain further insights into the structural basis of H2B.W2 function, we determined the cryo-EM structure of the H2B.W2 nucleosome (PDB: 9JC6) core particle (NCP) using 601 DNA (Supplementary Figure 5 and 6) and compared it with the structures of H2B (PDB: 8JBX) and H2B.W1 (PDB: 8CJJ) NCPs(5). A total of 264,491 particles were selected from micrographs of H2B.W2 NCPs and classified into several distinct groups (Supplementary Figure S6). From these, we chose 127,551 particles of homogeneous H2B.W2 NCPs, resulting in a final cryo-EM map with a resolution of 3.34 Å (Figure 2A, Supplementary Table 1 and Supplementary Figure 6).

**Figure 2.**
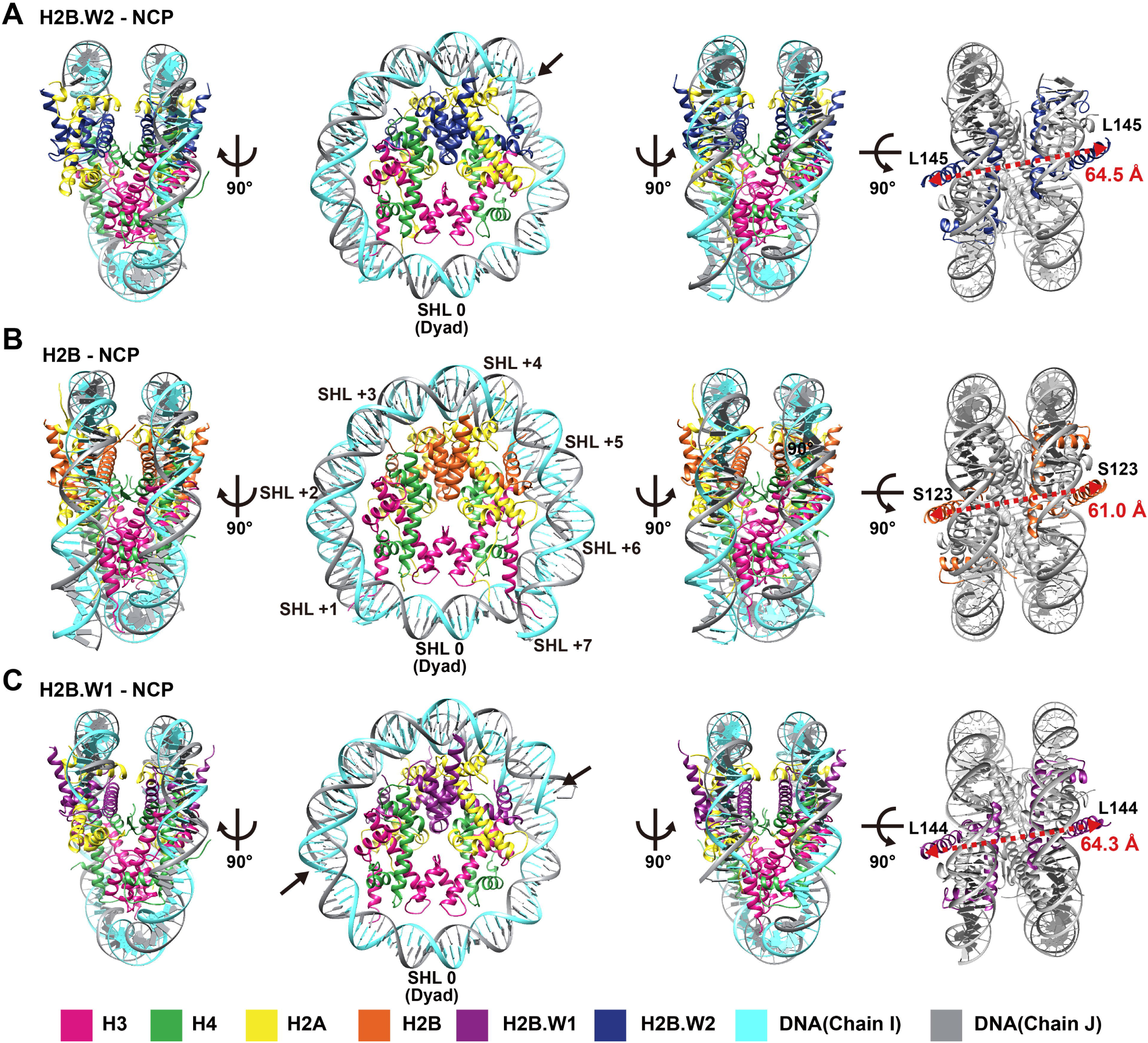
Cryo-EM structure of nucleosomes containing human H2B.W2. (**A-C**) Atomic models of (**A**) H2B.W2-NCP, (**B**) H2B-NCP, and (**C**) H2B.W1-NCP, in disc (middle) and gyre (left and right) views. The right most panel shows that distance changes between H2BS123-H2B’S123 in the H2B-NCP, H2B.W2L145-H2B.W2’L145 in the H2B.W2-NCP, and H2B.W1L144-H2B.W1’L144 in the H2B.W1-NCP.

Our structural analysis indicated that the H2B.W2-NCP is similar to the H2B-NCP in overall structure (Figure 2A, 2B and Supplementary Figure 7A), as well as the H2B.W2 and H2B structures (Figure 3A). However, we observed distinct conformational and structural rearrangements, including a flexible DNA end and an increased width of the nucleosome (Figure 2A, 2B and Supplementary Figure 7A). Specifically, in the H2B-NCP, we could resolve 147 bp of nucleosomal DNA, but the last 25 bp at the 3’ end of the 601 sequence of the H2B.W2 nucleosomal DNA remained unresolved. Furthermore, the nucleosome width— in terms of the distance from H2BS123 to H2B’S123, or from H2B.W2L145 to H2B.W2’L145—increased by 3.5 Å, from 61.0 Å in H2B-NCP to 64.5 Å in H2B.W2-NCP (Figure 2A and 2B).

**Figure 3.**
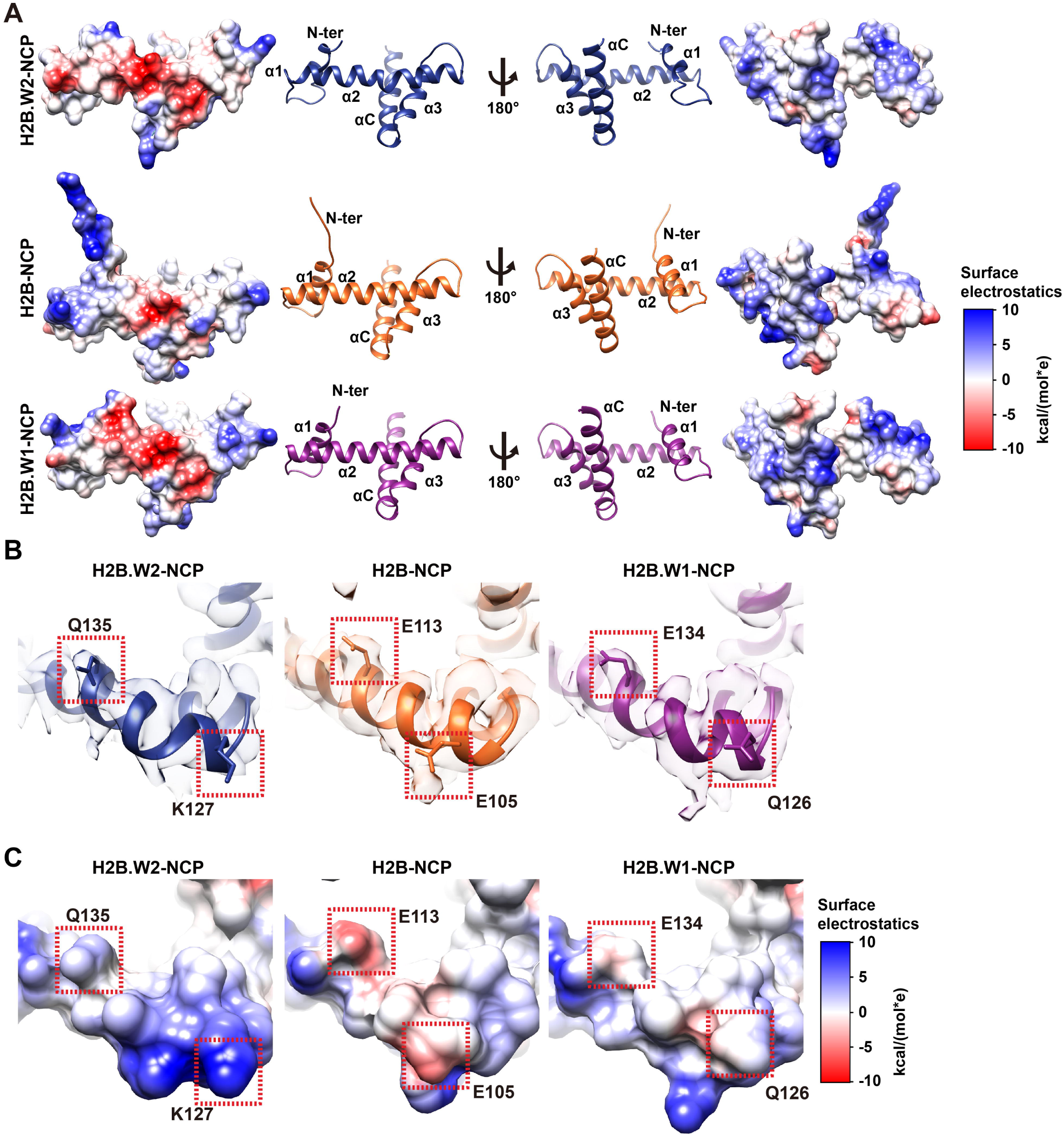
H2B.W2 increases the negative charges including the acidic patch region. (**A**) Comparison of the overall structure of H2B.W2, H2B, and H2B.W1. The red and blue indicate negative and positive, respectively, and the potential display levels are between -10 and 10 kcal/(mol*e)). (**B)** Close-up views of the acidic patch region of H2B.W2 (Left), H2B (Middle) and H2B.W1 (Right). EM density map superimposed with the atomic model of the acidic patch region. The regions bounded by the dotted red squares are those regions of the acidic patch that are different, and the same in (**C**). (**C**) Comparison of the surface electrostatic charges in the nucleosome acidic patch region in H2B.W2-NCP (Left), H2B-NCP (Middle) and H2B.W1-NCP (Right). The potential display levels are between -10 and 10 kcal/(mol*e)).

A similar increase in nucleosome width was also observed in the H2B.W1-NCP; however, the H2B.W1-NCP was missing 11 bp at one end and 18 bp at the other compared to the H2B-NCP (Figure 2C)(5). The similar asymmetry flexibility of both end of DNA was also observed in the CENPA-NCP(21). Whereas the flexible DNA of one side in the H2B.W2-NCP is around 25 bp, which is resemble to the hexasome structure(22). The hexasome has an additional ∼35 bp of unwrapped DNA, causes H3-H4 exposure, and enhances the activity of chromatin remodeler INO80(23,24). Therefore, this H2B.W2 destabilized nucleosome may also provide unique mechanism.

H2A-R77 and DNA form the minor groove anchor, and this anchor is critical for nucleosome stability(25). The DNA around one copy of H2A-R77 in the H2B.W2-NCP was invisible (Supplementary Figure 7B). This structural change was only observed on one side, and H2A’-R77 in the H2B.W2-NCP was inserted into a minor grove of DNA on the other side (Supplementary Figure 7B). Therefore, the H2B.W2 destabilizes the nucleosome around the dimer and DNA interaction regions, which is similar to the H2B.W1-NCP with a lower stability compared to H2B-NCP. However, the dimer DNA interaction difference between H2B.W1 and H2B.W2 nucleosomes indicates difference at other regions within these two variants also contribute to the nucleosome structure change.

To further confirm the length of DNA protected by the H2B.W2-NCP, we performed an MNase sensitivity assay. As expected, we observed a ∼150 bp band after MNase digestion of the H2B-NCP, consistent with previous studies (Figure 4A)(26). In contrast, digestion of the H2B.W2-NCP initially produced a band around 140 bp, which then shifted to a primary band around 120 bp, with even shorter fragments appearing over time (Figure 4A). This result aligns with the DNA length observed in our cryo-EM analysis of the H2B.W2-NCP. Additionally, based on the cryo-EM structure, the extended N-terminal tail may contribute to the protection of DNA fragment size. Therefore, we made an artificial construct swapping the N-terminal tail from H2B.W2 to H2B (i.e. H2B.W2-s.H2B-Nter). Although the overall protection size of the H2B.W2-sH2B-Nter-NCP was similar to that of the wild-type H2B.W2-NCP, the 120 bp band was more persistent in the chimeric nucleosome (Supplementary Figure 8). These findings suggest that while the extended H2B.W2 N-terminal tail contributes to the destabilization of nucleosomal DNA ends, other regions within H2B.W2 likely play a more dominant role in altering DNA-histone interactions and overall nucleosome stability.

**Figure 4.**
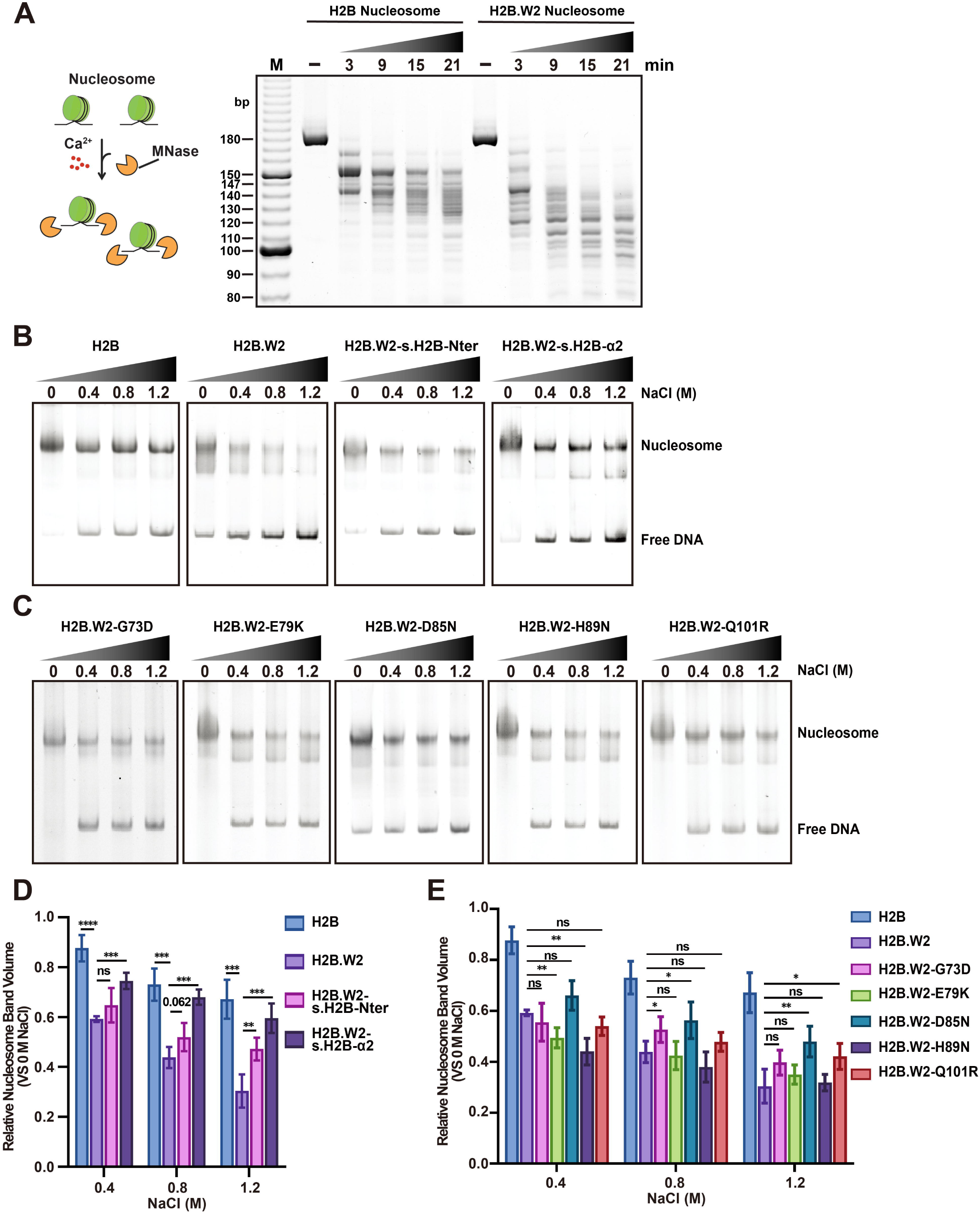
The stability of the H2B.W2 nucleosome is decreased and the H2B.W2 forms a smaller nucleosome. (**A**) Representative 12% 0.5x TBE gel image of H2B and H2B.W2 nucleosome samples after MNase digestion. (**B**) Nucleosome stability assay of the H2B, the H2B.W2 and the domain swapping nucleosomes. Polyacrylamide gels of the nucleosome after they were treated with different ionic concentrations (i.e., 0, 0.4, 0.8, or 1.2 M) of NaCl. (**C**) 0.5xTBE 6% polyacrylamide gels of the H2B.W2-G73D, H2B.W2-E79K, H2B.W2-D85N, H2B.W2-H98N, and H2B.W2-Q101R nucleosomes after they were treated with different ionic concentrations (i.e., 0, 0.4, 0.8, or 1.2 M) of NaCl. (**D**) Quantification of the nucleosome band in the gels from (**B**). The data (n=3) were compared using a Student t-test (two-tail, equal and unequal variances) and the asterisks indicate statistically significant differences at p < 0.01 (**), p < 0.001 (**\*\*\***), or p < 0.0001 (**\*\*\*\***), ns represents no significant statistical difference. (**E**) Quantification of the nucleosome band in the gels from (**C**). The data (n=3) were compared using a Student t-test (two-tail, equal and unequal variances) and the asterisks indicate statistically significant differences at p < 0.05 (*), p < 0.01 (**),, ns represents no significant statistical difference.

In addition to the observed differences in mononucleosome stability, our cryo-EM structure revealed alterations in the acidic patch of the H2B.W2 nucleosome. Specifically, two acidic patch amino acids in H2B (E105 and E113) are replaced with a positive residue (K127) and a neutral residue (Q135) in H2B.W2, while H2B.W1 only has one amino acid changed (Figure 3B, C and Supplementary Figure 1). The acidic patch, formed by six H2A and two H2B amino acids, is well known to play a role in chromatin-level interactions, influencing nucleosome-nucleosome interactions, and the chromatin remodelling complex interactions(27). These alternations in H2B.W2 could potentially influence higher-order chromatin structure and dynamics by modulating interactions with neighboring nucleosomes or altering the binding of chromatin remodeling factors.

### The negatively charged N-terminal tail, and D85 and Q101 in the α2 helix underlie the H2B.W2-nucleosome destabilization

To understand which domains of H2B.W2 contribute to its destabilizing effect, we systematically swapped each domain with its counterpart from H2B and examined their stability under high salt environments (Figure 4B, D and Supplementary Figure 9). These experiments revealed that, in addition to some contribution from the N-terminal tail, the α2 helix region (H2B.W2 79-105) is the major determinant of H2B.W2-NCP stability (Figure 4B, D and Supplementary Figure 9). This result is consistent with a recent solid-state NMR study, which found that part of the H2B α2 helix region positioned near the H4 α3 helix exhibits enhanced structural dynamics and is likely engaged in the coordination of octamer-DNA interactions(28). Given that the α2 helix region of H2B.W2 has more negative charge residues compared to H2B (Figure 1A), we hypothesized that these charges might be responsible for the H2B.W2-NCP destabilization. To test this hypothesis, we created a series of charge-altering mutants within the L1 loop and α2 helix regions (G73D, E79K, D85N, H89N, Q101R; Figure 4C, E). Salt stability assays of these mutants showed that neutralizing the charge of either D85 or Q101 (H2B.W2 D85N and H2B.W2 Q101R) significantly increased nucleosome stability compared to the wild-type H2B.W2 (Figure 4C, E).

Further examining the cryo-EM structure of the H2B.W2-NCP, we found that D85, a negatively charged residue, faces the N-terminal tail of H2B.W2 (Supplementary Figure 10A). Substituting this residue with a similar sized but neutral Asn residue (i.e. D85N) enhanced the stability of the nucleosome possibly by increasing the overall positive charge of the core octamer and thus strengthening the DNA-octamer interactions (Supplementary Figure 10A). This implies that the D85 residue of H2B.W2 likely contributes to nucleosome destabilization via enhanced charge-repulsion. Furthermore, Q101 is positioned within the core histone octamer, and while the additional positive charge from the R mutation might help improve nucleosome stability, the detailed mechanism by which Q101 contributes to the overall lower stability of the H2B.W2-NCP compared to canonical H2B remains unclear (Supplementary Fig. 10B).

### H2B.W2 destabilizes the nucleosome by ∼35% along the interaction regions between histone H2A-H2B dimers and DNA

To quantify the nucleosome destabilization effects of H2B.W2, we performed single-molecule optical tweezers experiments. Optical tweezers experiments allow us for precise force measurement to extract the strength of histone-DNA interaction. Nucleosomes were attached to biotin- and digoxigenin-labeled DNA handles (Figure 5A). When the applied force was increased in the nucleosome pulling assay, we observed two distinct unwrapping events: a lower-force event (outer rip) and a higher-force event (inner rip)(29) (Figure 5B). The outer rip corresponds to the unwrapping of the outer DNA turn, releasing ∼60-70 bp of DNA, while the inner rip reflects the unwrapping of the inner DNA turn, releasing ∼70-80 bp. Note that the unwrapping force and the size of the outer rip were slightly larger than those reported in a recent single-molecule study(30) (Supplementary Table 2). This discrepancy may be attributed to the presence of linker histone H1.4, as well as variations in salt concentration and the type of salt used.

**Figure 5.**
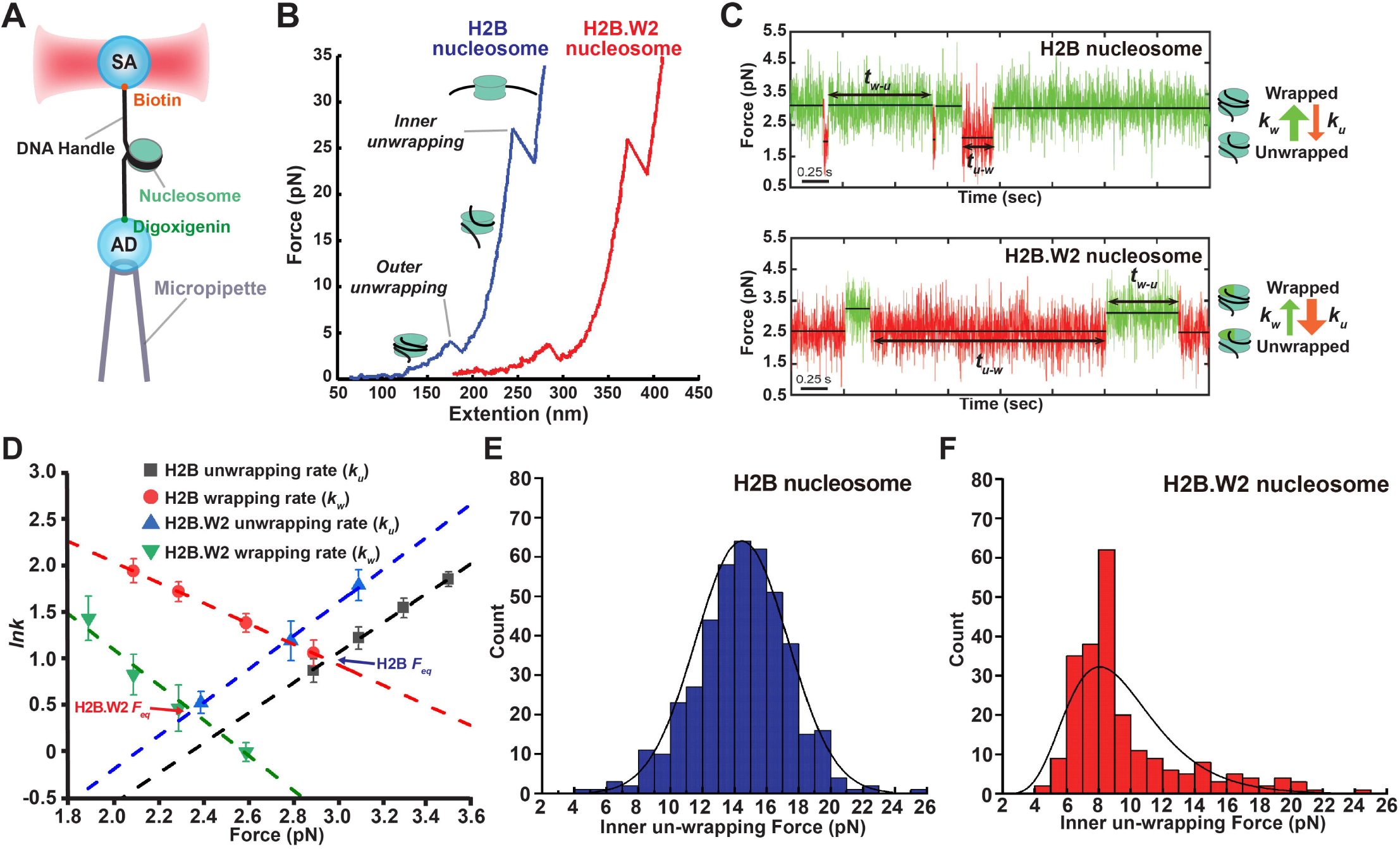
The H2B.W2 nucleosome destabilizes the interaction between DNA and H2A-H2B.W2 by ∼35% in single molecule optical tweezers assays. (**A**) Schematic to show the optical tweezers setup used in the nucleosome studies. SA represents a streptavidin coated bead and AD represents an anti-digoxigenin coated bead. (**B**) Comparison of two typical force-extension curves obtained for canonical H2B (blue) and H2B.W2 (red) nucleosomes. (**C**) The representative wrapping and unwrapping events of the canonical H2B and H2B.W2 nucleosomes in the outer hopping assay. (**D**) Quantification of the unwrapping and rewrapping rates under different forces. The dashed lines represent lines of best fit. Data indicate the mean ± SEM. (**e** and **f**) Distribution inner unwrapping force for the H2B and the H2B.W2 nucleosomes. (**E**) Inner unwrapping force observed for the H2B nucleosome, main population (10-18 pN). (**F**) Inner unwrapping force observed for the H2B.W2 nucleosome, main population (6-10 pN).

To quantify the effect of H2B.W2 on the outer rips, we performed a single-molecule optical tweezers nucleosome hopping assay at 5 mM NaCl. In this assay, we held a nucleosome tether at a constant trap position under varying force ranges to observe transitions between the wrapped and unwrapped state (Figure 5C, D). In the H2B.W2-NCP, the lifetime of the nucleosome wrapped stage *t_w-u_* was decreased, and the lifetime of the nucleosome unwrapped state *t_u-w_* was increased compared to the H2B-NCP (Figure 5C, D). Then, we extracted the equilibrium force (*F_eq_*), the force at which *t_w-u_* equates to *t_u-w_*, was reduced from 2.97 pN in the H2B-NCP to 2.36 pN in the H2B.W2-NCP (Figure 5C, D, and Supplementary Table 2). We then calculated the free energy cost of the outer rip at 0 pN (Δ*G°*) to be 20.46 kJ/mol for the H2B-NCP, whereas it was significantly reduced to 13.22 kJ/mol for the H2B.W2-NCP(13–15). This 35% reduction in the energy barrier for the outer rip in the presence of H2B.W2 is similar to the 37% reduction observed in the presence of H2B.W1(5).

Average inner rip forces were at around ∼15 pN and ∼7 pN in the H2B-NCP and H2B.W2-NCP, respectively, in presence of 300 mM NaCl (Figure 5E, F). These results are consistent with our MNase and salt stability assay data, suggesting that weakened interactions between the H2A-H2B.W2 dimer and the rest of the nucleosome promote dissociation of the dimer. This dissociation might lead to the formation of a tetrasome, which could explain the lower inner unwrapping force (∼ 7 pN) observed in H2B.W2-NCP (Figure 5E, F). These results indicate a substantial impact of H2BW2 on nucleosome dynamics.

### H2B.W2 disrupts chromatin compaction through G73 in the L1 loop

Beyond the mononucleosome level, chromatin structure is the primary biologically functional form under *in vivo* condition, and nucleosomal DNA end interactions and linker DNA length are known to influence chromatin folding(31,32). We investigated whether the increased DNA ends flexibility in H2B.W2 nucleosomes impacts higher-order structure. We compared the stacking dynamics of H2B and H2B.W2 tri-nucleosomes, representing the minimal unit of chromatin, across varying Mg^2+^ concentrations (Figure 6 and Supplementary Figure 11)(16,33). The H2B tri-nucleosome adopted a more compact structure with increasing Mg^2+^ concentration (Supplementary Fig. 11A, C), which is consistent with previous reports(16,33). A similar trend was observed for the H2B.W2 tri-nucleosome, however, the initial FRET signal was much lower in the H2B.W2 condition, indicating a less compact chromatin conformation at low Mg^2+^ concentrations (Supplementary Figure 11). This relative openness is maintained across all Mg^2+^ concentrations tested, which encompassed the physiological Mg^2+^ concentration at around 1 mM (Supplementary Figure 11). Notably, a crystal structure of a 12-nucleosomal fiber revealed that H2BD51, the canonical H2B residue equivalent to G73 in H2B.W2, within the L1 loop interacts with H2AR71 in neighboring nucleosomes(34).

**Figure 6.**
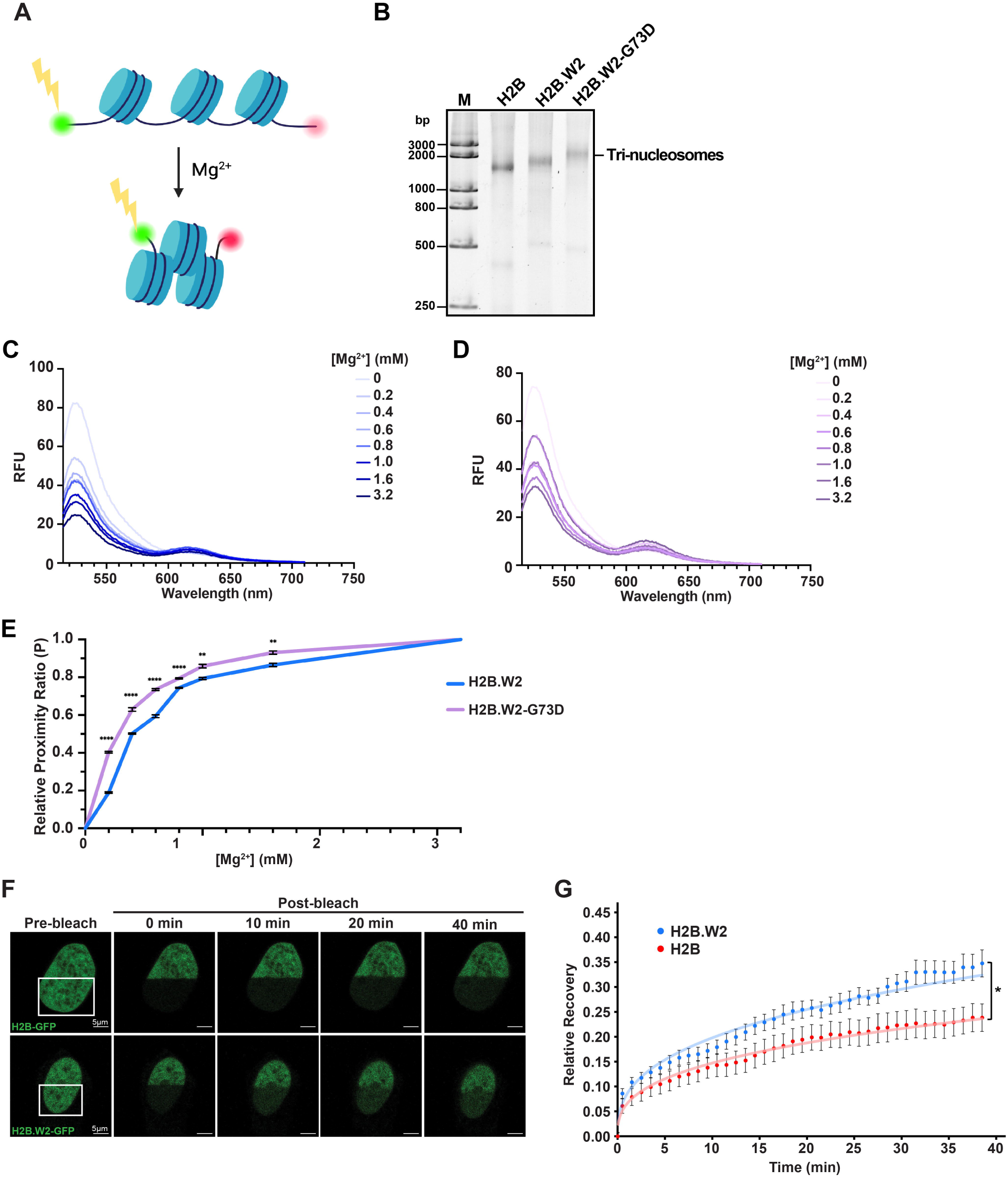
H2B.W2G73 plays an important role in opening up the chromatin structure composed of H2B.W2. (**A**) The schematic diagram of tri-nucleosome array condensation FRET assay. (**B**) Representative native-PAGE of the loaded H2B, H2B.W2, and H2B.W2-G73D tri-nucleosomes. (**C** and **D**) Representative emission spectrums of (**C**) H2B.W2 tri-nucleosome, and (**D**) H2B.W2-G73D tri-nucleosome from 515 nm to 710 nm upon excitation at 488 nm exposed to the indicated amounts of Mg^2+^ ions. (**E**) The relationship between the tri-nucleosome array FRET proximity ratios (P) and the Mg^2+^ ion concentrations using different Mg^2+^ concentrations. Each datapoint represents the mean proximity ratio ± 1 S.D. (n=3 for all conditions tested). The data were compared with a Student t-test and the asterisks indicate statistically significant differences at p < 0.01(**), or p < 0.0001 (****). (**F**) Representative fluorescence images of the HeLa cell nuclei expressing either H2B-GFP or H2B.W2-GFP pre-bleach and 0, 10, 20, 40 min post-bleach. (**G**) Line graph showing the relative fluorescence signal recovery after photo-bleaching. The error bars indicate the SEM and the p-values were calculated with the Student t-test (two tail, unequal variance) (n = 13 for H2B.W2 & n = 9 for H2B (H2B dataset is from (5)).

We hypothesized that H2B.W2G73 may change the interaction with H2AR71, thus changing the chromatin structure. The decreased chromatin compaction of H2B.W2 tri-nucleosome is partially reverted by the H2B.W2-G73D mutation (Figure 6C, D, E). These results indicate that H2B.W2 not only destabilizes individual nucleosomes but also disrupts higher-order chromatin structure, with G73 playing a key role in this destabilization at the chromatin level.

To further validate the destabilizing effect of H2B.W2 on nucleosomes and chromatin within a cellular context, we performed fluorescence recovery after photobleaching (FRAP) experiments using EGFP-tagged H2B or H2B.W2 (Figure 6F, G). As expected, EGFP-tagged H2B or H2B.W2 were both shown to be localized in the nucleus (Figure 6F). As previously reported, the EGFP fluorescence recovery of canonical H2B was very slow because canonical histones are stably bound to chromatin and thus virtually immobile (Figure 6F, G)(35–37). In contrast, the H2B.W2-EGFP signal showed a significantly faster recovery. These results suggest that H2B.W2 is in fact more mobile and diffusive compared to canonical H2B in cells.

## Discussion

Our study reveals that H2B.W2, a testis-specific histone variant, exhibits unique structural and dynamic properties that distinguish it from canonical H2B and its paralog H2B.W1. Specifically, we found that H2B.W2 destabilizes nucleosomes, disrupts chromatin compaction, and exhibits increased mobility within the nucleus. These findings suggest a specialized role for H2B.W2 in regulating chromatin structure and accessibility during spermatogenesis. The timing of H2B.W2 expression in spermatogenesis is distinct from that of H2B.C1, which is a well-studied testis-specific variant of histone H2B. H2B.C1 is expressed in the later stages of spermatogenesis, particularly in the spermatids, and it is known to be distributed evenly across whole chromosomes(38). The histone variant evolutionarily closest to H2B.W2 is H2B.W1; it is expressed slightly earlier than H2B.W2 but there is some overlap in the expression stages. Single-cell RNA sequence analysis showed that ∼40% of the cells that express H2B.W1, also express H2B.W2 (Figure 1). H2B.W1-NCP and H2B.W2-NCP have similar nucleosome destabilizing effects and are expressed in the early stages of spermatogenesis. Although it is still not clear whether disruption of H2B.W2 function might affect human spermatogenesis, disruptions of H2B.W1 gene (i.e., two SNPs of the H2B.W1 at -9C>T and 368A>G) are known to be related to infertility(8,9). Therefore, it is possible that mutations of H2B.W2 will also be associated to infertility. Further studies would be helpful to understand how H2B.W1 and H2B.W2 function in spermatogenesis.

The significant decrease in H2B.W2-NCP stability compared to canonical nucleosomes suggests a more dynamic role for this variant. Based on the amino acid sequence alignment and the domain swapping salt stability assay, the H2B.W2 N-terminal tail and the α2 helix significantly contributed to the nucleosome destabilization. The N terminus of H2B.W2 contains much more negatively charged amino acid residues than that of H2B, and slightly more than that of H2B.W1. Previous studies have shown that the H2B N-terminus can adopt different orientations within the nucleosome, influencing DNA-histone interactions(39). We speculate that the excess negative charge in H2B.W2 disrupts these interactions through enhanced electrostatic repulsion with DNA. This is supported by our cryo-EM structure, which reveals increased flexibility in the H2B.W2 N-terminal region.

The observation that H2B.W2-NCP protected a shorter DNA region, potentially indicative of hexasome formation, further supports its role in creating a more open chromatin conformation. While we cannot exclude the possibility of hexasome formation, which has a quite low chance since the two copies of H2A/H2B.W1 dimer density map resolution have no huge difference, both asymmetric unwrapping and hexasome structures represent weaker conformations that could facilitate transcription. This is particularly relevant for H2B.W2, as its destabilization might be crucial for upregulating genes expressed during early spermatogenesis.

In addition to its impact on nucleosome stability, H2B.W2 also influences for the chromatin stability. G73, located in the L1 loop of H2B.W2, appears to play a role in chromatin compaction, potentially by reducing the attraction with H2AR71. This suggests that while certain regions of H2B.W2 contribute to nucleosome destabilization, other parts may be involved in modulating higher-order chromatin structure. Furthermore, our FRAP assay revealed a faster signal recovery for EGFP-tagged H2B.W2 compared to EGFP-tagged H2B, suggesting increased mobility of H2B.W2 within the nucleus. This observation aligns with recent findings highlighting the importance of positive charges in histone tails for stable nucleosome association(40). Including the changes in the acidic patch region, these results suggest that the altered charge distribution in H2B.W2 not only destabilizes its nucleosome structure but also makes it more accessible to chromatin remodelers and histone chaperones or promote the recruitment of spermatogenesis-specific factors. This instability might be helpful during early spermatogenesis, where rapid chromatin remodeling is required for meiosis and later histone to protamine replacements.

This increased accessibility, coupled with the potential for hexasome formation, raises intriguing questions about the localization and function of H2B.W2 during spermatogenesis. While canonical histones typically exhibit a more widespread distribution, unique properties of H2B.W2 might target it to specific genomic regions, as hinted by the uneven and dotted staining patterns observed in H2B.W2-positive spermatocytes (Figures 1A, B and Supplementary Figure 3). Interestingly, H2B.W1 has been found to localize at telomeric regions in the V79 cell line(6). Transiently expressed H2B.W2 in HeLa cells also localized to sub-telomeric and telomeric regions (data not shown), it is possible that the function of H2B.W2 may be similar to that of H2B.W1. Since spermatogenic cells are different from cultured cell lines, both H2B.W2 and H2B.W1 may be localized at different locus in male germ cells. Though, it is interesting to investigate whether H2B.W2 and H2B.W1 are localized at telomeric regions during spermatogenesis and is involved in telomere maintenance. Further investigation of the precise mechanisms by which H2B.W2 destabilizes chromatin structure within the context of native genomic sequences will be essential for fully understanding the complexities of spermatogenesis and male fertility.

## Acknowledgements

This work was supported grants from the Hong Kong Innovation & Technology Commission Innovation and Technology Fund [MHP/033/20], the National Natural Science Foundation of China [32170548] awarded to T.I., Shenzhen Science and Technology Innovation Commission [JCYJ20200109140201722] to YY, and grants from the Research Grants Council of the Hong Kong SAR [C6036-21GF] awarded to Y.Z. and T.I. and [16103620, T13-602/21N] to YY. EM dataset was collected at the Biological Cryo-EM Center, generously supported by a donation from the Lo Kwee Seong Foundation, at HKUST.

