## supplemental document for "H2B.W2, a Spermatocytes-specific Histone Variant, disrupts nucleosome stability and reduces chromatin compaction"

### Supplemental Material and methods

#### Expression and purification of recombinant proteins

Each of the human canonical histone H2A, H2B, H3 and H4 was cloned into pET11A, and their expression and purification were conducted as previously described(1). His-SUMO-H2B.W2 was cloned into the pET28A vector and expressed in *E. coli* BL21(DE3). The recombinant protein was then purified through a Ni-NTA resin (Qiagen). The His-SUMO tag was removed using a digestion buffer comprising 1x PBS containing 150 mM NaCl, 1% Triton X-100 and the *PreScission* protease (GE Healthcare) at 4°C for 24 h. After the tag removal, the digested elution sample was adjusted to 0.4 N HCl and centrifuged at 36,000 g, at 4°C for 20 mins to remove any non-histone proteins. Recombinant H2BW2 was then concentrated by acetone precipitation.

#### Energy barrier extraction from nucleosome hoping events

The free energy $\Delta G^0$ was calculated with the following the formula: $\Delta G^0 = F_{eq}\Delta x - \Delta G_{strech} - k_{BT} ln(k_{eq})$, in which, $\Delta G_{stretch}$ is the energy needed to stretch the free DNA template to the equilibrium force ($F_{eq}$) and the equilibrium rate ($k_{eq}$) is defined as the stage that $k_w$ is equal to $k_u$. $\Delta x$ is the average DNA extension change in the outer pulling assay, $k_{BT}$ is the constant value generated from the Boltzmann constant ($kB$) and the temperature ($T$) (2-4) .

**Supplementary Figure 1 Amino acid sequence alignment of *Hs*H2B, *Hs*H2B.W1 and *Hs*H2B.W2**

The N- and C-terminal tail regions are indicated by black lines and the histone fold domains are indicated by the orange rectangles. The negatively and positively charged amino acids are shown in red and cyan, respectively. Identical positions are highlighted in dark green, whereas similar positions are highlighted in light green. Black arrow indicates the site of point mutation. The wavy underlined sequence indicates the peptide epitopes used for generating the *Hs*H2B.W2 antibody.

**Supplementary Figure 2 Validation of human H2B.W2 antibody**

(**A-C**) Representative image resulted from western-blot analysis of recombinant proteins (**A**) or HEK293T cells without or with overexpression of H2B.W2-3xFlag (**B** and **C**) using rabbit-anti human H2B.W2 antibody (**A** and **C**) and mouse anti-Flag antibody (**B**).

**Supplementary Figure 3 Additional immunofluorescence images of the human testis sections showing the localization of H2B.W2 and γ-H2A.X**

Three representative human testis sections were dual-immunolabeled with anti-H2B.W2 antibody (red) and anti-phospho-histone H2A.X (γ-H2A.X; green) antibodies. DNA was counterstained with Hoechst 33342 (blue). Scale bar = 50 μm.

**Supplementary Figure 4 Single-cell RNA expression patterns of the spermatogenesis marker genes, (STRA8 and SPO11), as well as H2B.C1, H2B.W1 and H2B.W2 during spermatogenesis, with UMAP analysis**

**Supplementary Figure 5 The H2B.W2-NCP preparation for cryo-EM image processing**

(**A**) SDS-PAGE of histone H2B octamer and H2B.W2 octamer. (**B** and **C**) Native-PAGE of the H2B-NCP (**B**) and H2B.W2-NCP (**C**) using the 147 bp Widom 601 nucleosome positioning sequence.

**Supplementary Figure 6 Representative micrographs and 2D classes and cryo-EM image processing of the H2B.W2-NCP**

(**A**) Representative micrographs of the H2B.W2-NCP. (**B**) Flowchart of cryo-EM data processing for the H2B.W2 nucleosome. In general, 41 2D classes were selected for the further analysis. (**C**) 246,491 particles were selected from a total of ~575,000 particles. These particles were further classified into several 3D classes. (**D**) Further analysis resulted in a final density map. (**E**) Fourier shell correlation curve of the H2B.W2NCP density map. The final resolution is 3.34 Å (purple) with golden standard. (**F**) The final 3D density map of the H2B.W2 nucleosome with the local resolution overlay.

**Supplementary Figure 7 Comparation of the H2B.W2, H2B, and H2B.W1 nucleosomes overall and the dimer-DNA interaction regions**

**(A)** EM density for H2B.W2 (Left), H2B (Middle) and H2B.W1 (Right) nucleosomes superimposed with the atomic models. **(B)** Comparison of the two copies (Upper and lower panels) of H2A/H2B dimer and DNA interaction among the H2B.W2 (Left), H2B (Middle) and H2B.W1 (Right) nucleosomes.

**Supplementary Figure 8 MNase digestion assay in the H2B.W2 and the H2B.W2-s.H2B-Nter nucleosomes**

Representative 12% 0.5x TBE gel image of H2B.W2 and H2B.W2-s.H2B-Nter nucleosome samples after MNase digestion.

**Supplementary Figure 9 The salt stability assay with swapping domain of H2B.W2**

(**A**) Representative salt stability assay gel of H2B.W2-$\Delta$0–21, H2B.W2-s.H2B-$\alpha$1, H2B.W2-s.H2B- $\alpha$3 and H2B.W2-s.H2B- $\alpha$C nucleosomes. (**B**) Relative nucleosome volume against 0 M NaCl among H2B, H2B.W2, H2B.W2-$\Delta$21, H2B.W2-s.H2B-$\alpha$1, H2B.W2-s.H2B-$\alpha$3 and H2B.W2-s.H2B-$\alpha$C nucleosomes.

**Supplementary Figure 10 The structural surface electrostatic properties change by D85N and Q101R mutations**

(**A**) Comparison of the surface electrostatic properties of D85 in H2B.W2-WT and N85 in the H2B.W2-D85N mutant. (**B**) Comparison of the surface electrostatic properties of Q101 in H2B.W2-WT and R101 in the H2B.W2-Q101R mutant. In both cases, red and blue indicate negative and positive charges, respectively, and the potential display levels range from -10 to 10 kcal/(mol*e)).

**Supplementary Figure 11 FRET assay of H2B, H2B.W2, and H2B.W2G73D tri-nucleosomes**

(**A**) Representative FRET fluorescence spectrums of H2B tri-nucleosomes in the presence of $Mg^{2+}$. (**B**) Representative FRET fluorescence spectrums of free DNA template in the presence of

$Mg^{2+}$. (**C**) Line graph of the relationship between the tri-nucleosome array FRET proximity ratios (P) and the $Mg^{2+}$ range. Each datapoint represents the mean P ± 1 S.D. (n = 3 for all conditions tested).

**Supplementary Table 1 Cryo-EM data collection, refinement, and validation statistics**

**Supplementary Table 2 Summary of wrapped rate and unwrapped rate in outer hopping assay.**
