## Supplementary figures and images for "H2B.W2, a Spermatocytes-specific Histone Variant, disrupts nucleosome stability and reduces chromatin compaction"

### Supplementary Figure 1

**Figure S1. Nguyen *et al.*,**

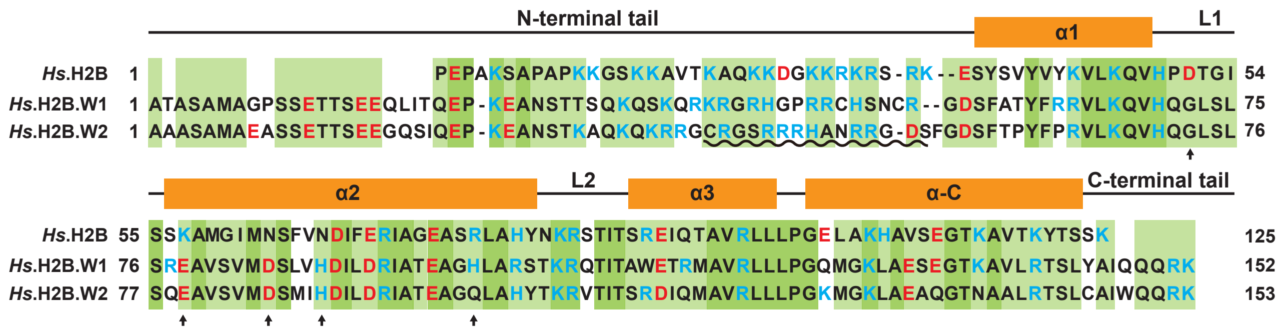

### Supplementary Figure 2

**Figure S2. Nguyen *et al.*,**

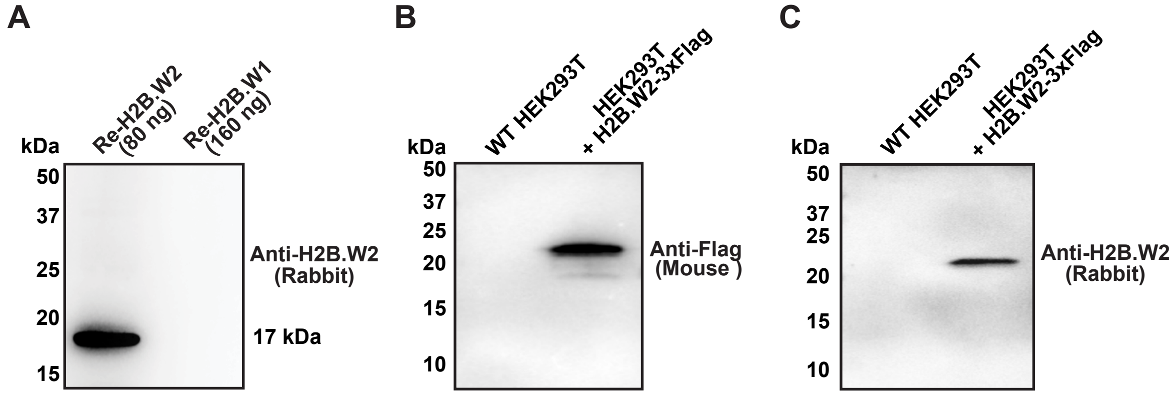

### Supplementary Figure 3

Figure S3. Nguyen *et al.*,

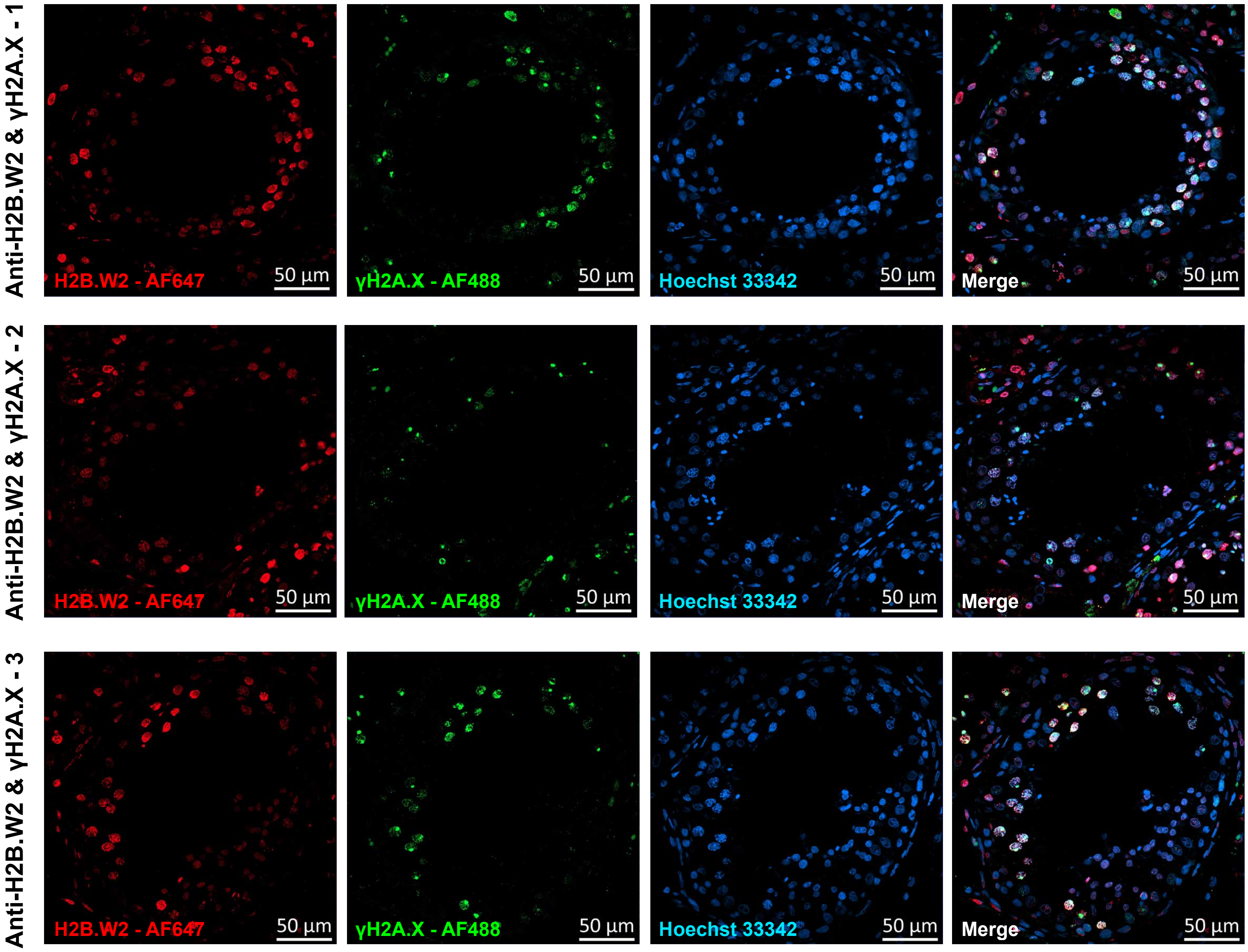

### Supplementary Figure 4

**Figure S4. Nguyen *et al.*,**

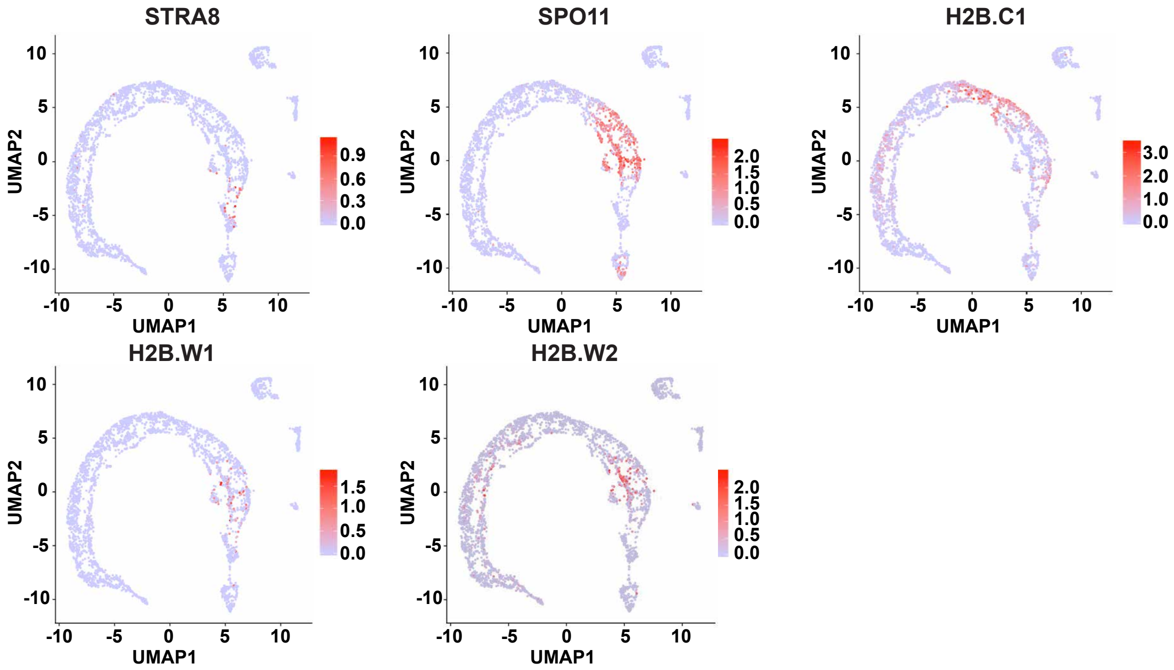

### Supplementary Figure 5

**Figure S5 Nguyen *et al.*,**

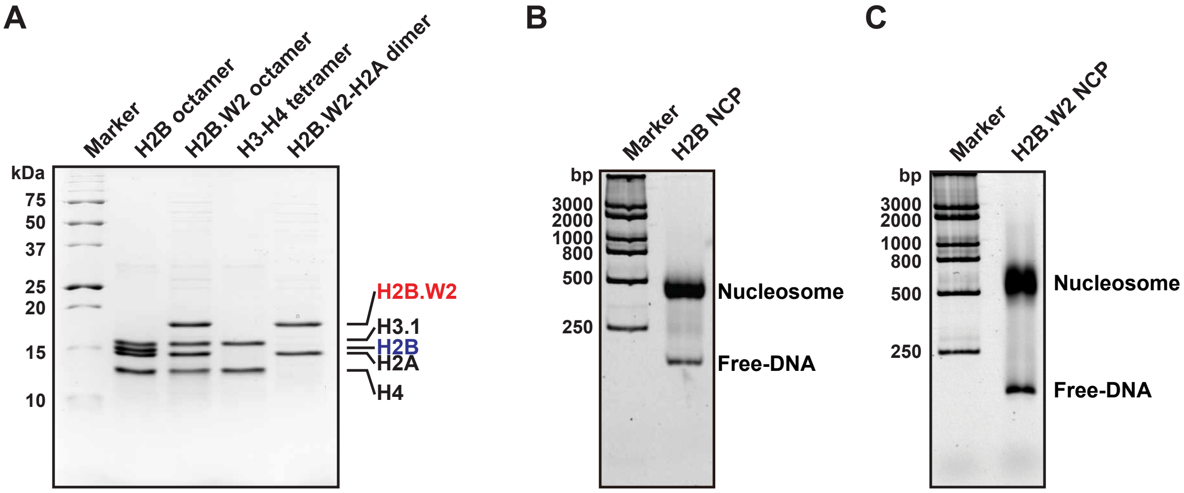

### Supplementary Figure 6

#### Figure S6. Nguyen *et al.*,

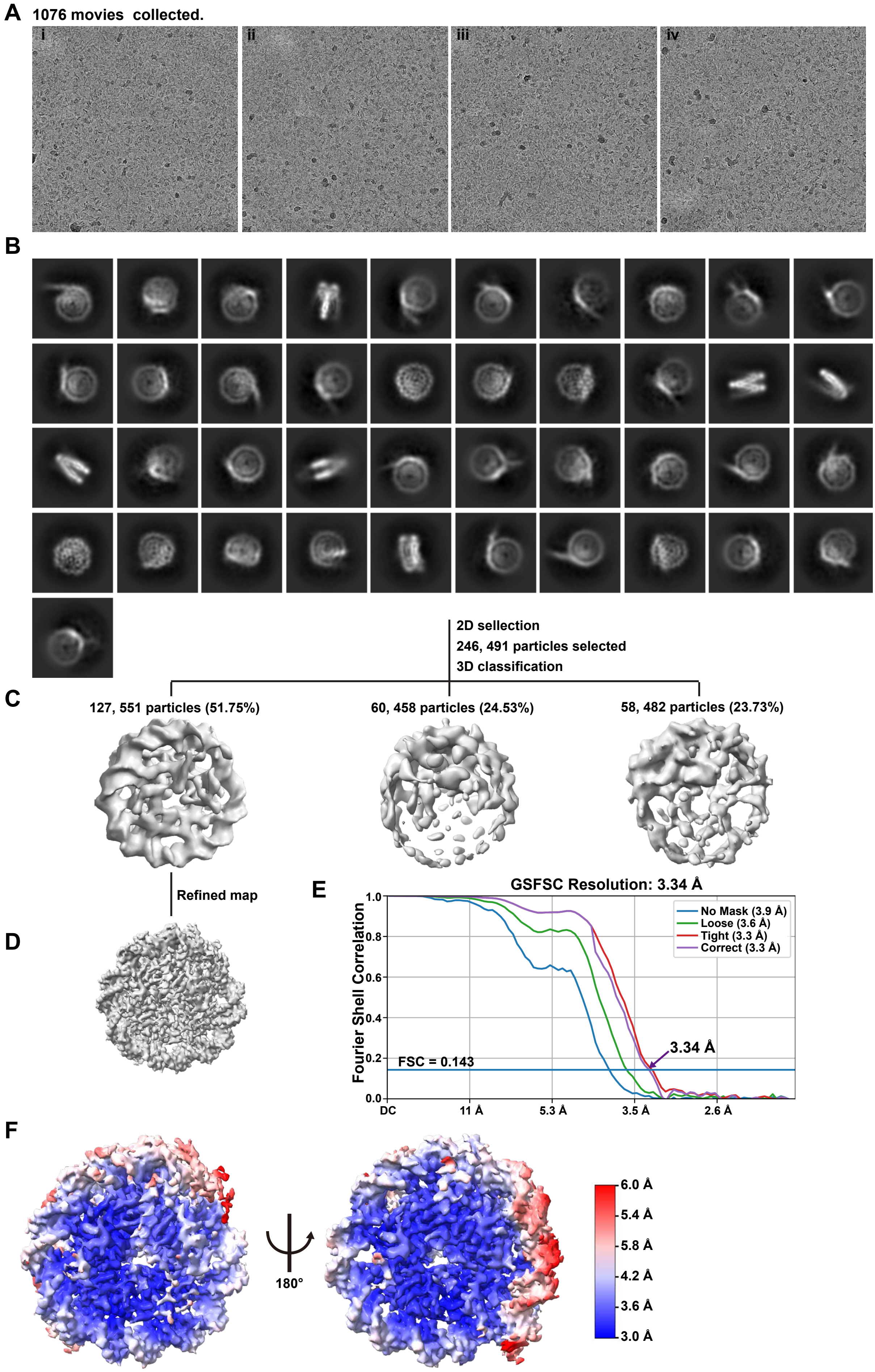

### Supplementary Figure 7

#### Figure S7 Nguyen *et al.*,

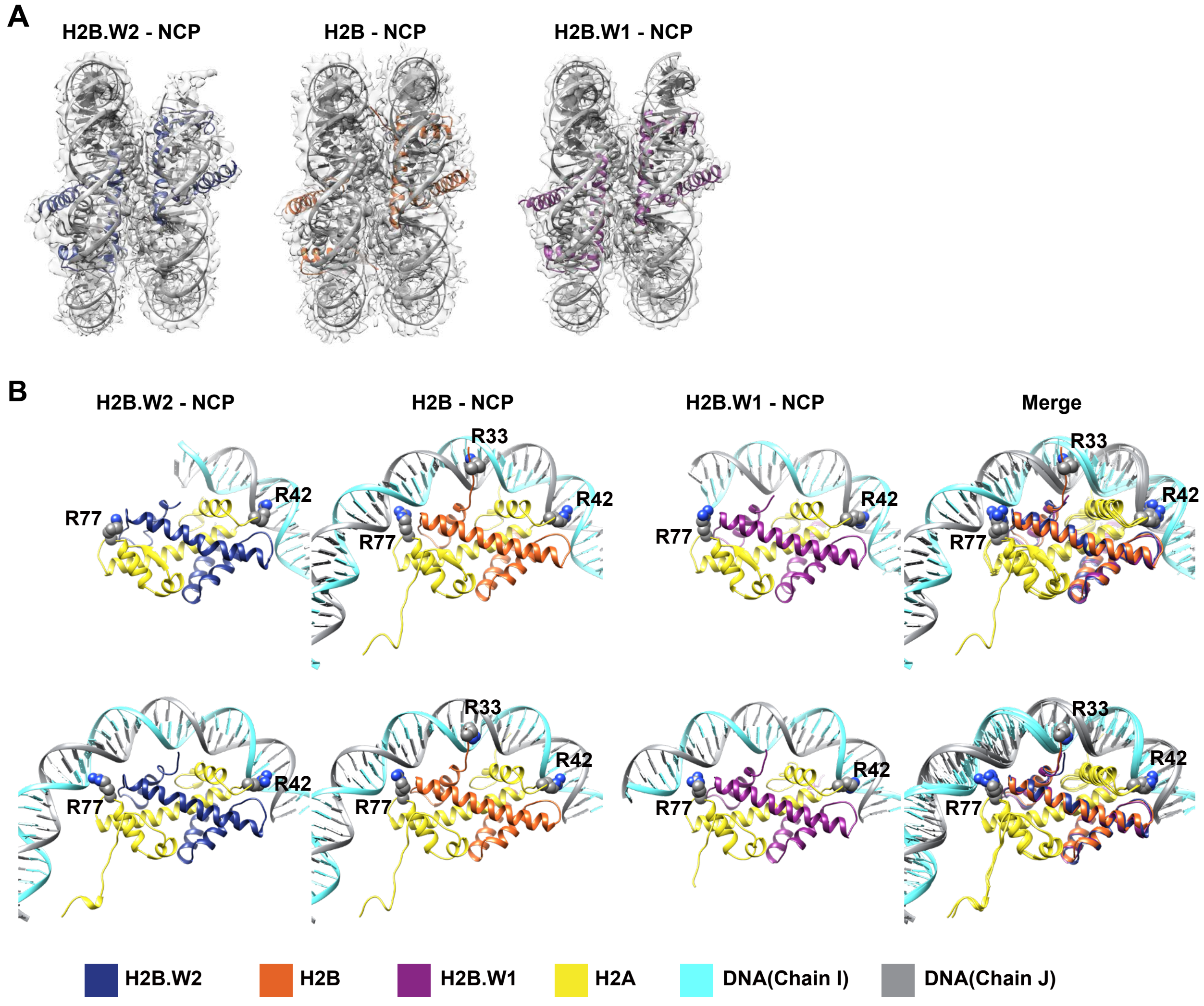

### Supplementary Figure 8

**Figure S8. Nguyen *et al*.,**

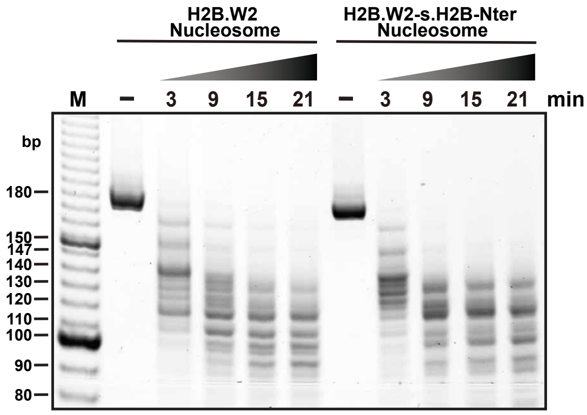

### Supplementary Figure 9

**Figure S9. Nguyen *et al.***

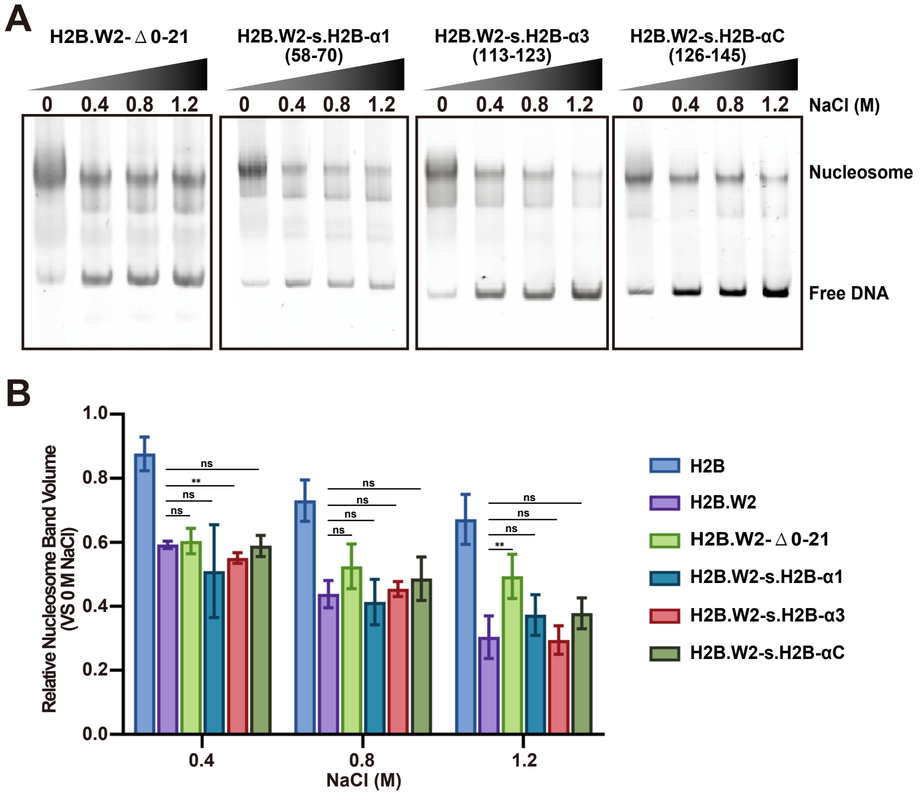

### Supplementary Figure 10

Figure S10 Nguyen *et al.,*

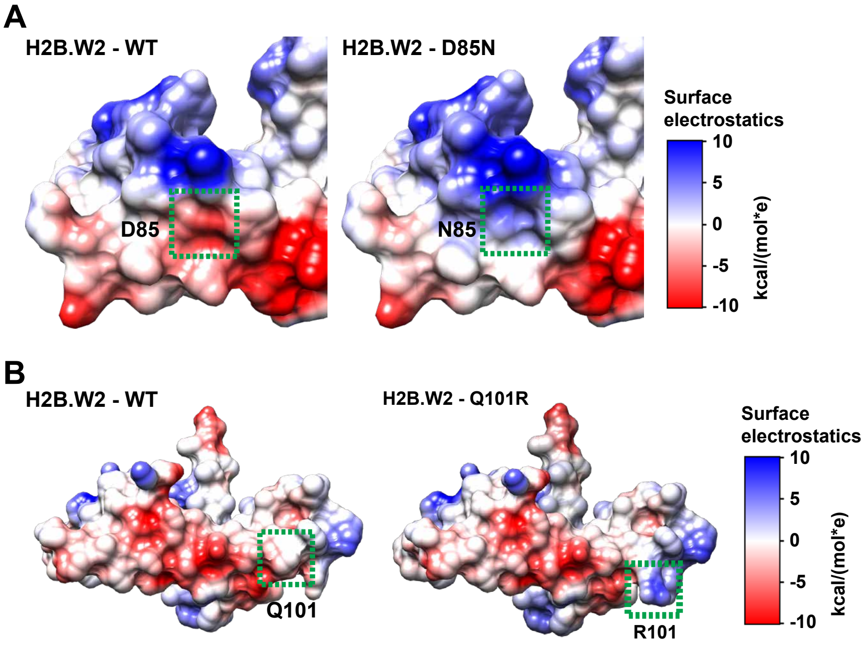

### Supplementary Figure 11

#### Figure S11. Nguyen *et al.*,

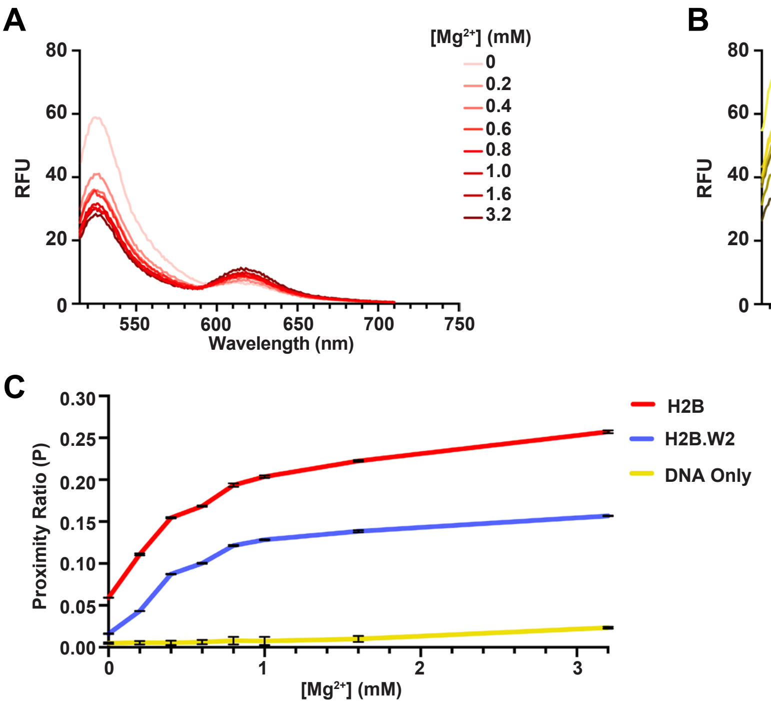
