## Supplementary material for "H2B.W2, a Spermatocytes-specific Histone Variant, disrupts nucleosome stability and reduces chromatin compaction": Table S1

**Table S1. Cryo-EM data collection, refinement, and validation statistics**

| | H2B.W2-NCP (EMD-61358) (PDB 9JC6) |
|---|---|
| **Data collection and processing** | |
| Number of frames collected | 40 |
| Defocus range (μm) | -0.25—-1.10 |
| Symmetry imposed | C1 |
| Micrographs recorded/used (no.) | 1076/622 |
| Initial particle images (no.) | 575,914 |
| Final particle images (no.) | 127, 551 |
| Final reconstruction package | cryoSPARC |
| Map resolution (Å) | 3.34 |
| FSC threshold | 0.143 |
| **Model Building** | |
| Software | Coot |
| Initial model used (PDB) | 3LZ0 |
| Refinement | Phenix |
| **Model composition** | |
| Protein residues | 706 |
| Nucleotide | 240 |
| **Validation** | |
| MolProbity score | 1.73 |
| Clashscore | 12.01 |
| Poor rotamers (%) | 0.00 |
| Bond lengths (Å) | 0.004 |
| Bond angles (˚) | 0.575 |
| **Ramachandran plot** | |
| Favored (%) | 97.25 |
| Allowed (%) | 2.75 |
| Disallowed (%) | 0 |
