## Supplementary material for "H2B.W2, a Spermatocytes-specific Histone Variant, disrupts nucleosome stability and reduces chromatin compaction": Table S2

**Table S2 Summary of wrapped rate and unwrapped rate in outer hopping assay.**

| $k_u$ | | | | $k_w$ | | | |
|---|---|---|---|---|---|---|---|
| H2B | | H2B.W2 | | H2B | | H2B.W2 | |
| Force (pN) | $k_u$ (n)* | Force (pN) | $k_u$ (n)* | Force (pN) | $k_w$ (n)* | Force (pN) | $k_w$ (n)* |
| 2.90 (2.50-2.98) | 2.4 ± 0.3 (111) | 2.40 (2.20-2.62) | 1.7 ± 0.2 (72) | 2.10 (1.90-2.15) | 7.0 ± 1.0 (81) | 1.90 (1.85-2.05) | 4.2 ± 1.0 (35) |
| 3.10 (2.98-3.18) | 3.4 ± 0.4 (133) | 2.70 (2.62-2.86) | 2.4 ± 0.6 (49) | 2.30 (2.15-2.38) | 5.6 ± 0.6 (143) | 2.10 (2.05-2.20) | 2.3 ± 0.5 (30) |
| 3.30 (3.18-3.45) | 4.7 ± 0.5 (157) | 2.90 (2.86-3.00) | 4.4 ± 2.0 (53) | 2.60 (2.38-2.78) | 4.0 ± 0.4 (240) | 2.30 (2.20-2.40) | 1.6 ± 0.4 (35) |
| 3.50 (3.45-3.63) | 6.4 ± 0.5 (123) | 3.10 (3.00-3.30) | 6.0 ± 1.0 (80) | 2.90 (2.78-2.95) | 2.9 ± 0.5 (107) | 2.60 (2.40-2.80) | 1.0 ± 0.1 (85) |

(n)* means the number of events happened at each force range.
